# A novel approach in heart-rate-variability analysis

**DOI:** 10.1101/2021.10.28.466234

**Authors:** András Búzás, Tamás Horváth, András Dér

## Abstract

Heart-rate variability (HRV), measured by the fluctuation of beat-to-beat intervals, has been growingly considered the most important hallmark of heart rate (HR) time series. HRV can be characterized by various statistical measures both in the time and frequency domains, or by nonlinear methods. During the past decades, an overwhelming amount of HRV data has been piled up in the research community, but the individual results are difficult to reconcile due to the different measuring conditions and the usually HR-dependent statistical HRV-parameters applied. Moreover, the precise HR-dependence of HRV parameters is not known. Using data gathered by a wearable sensor of combined heart-rate and actigraphy modalities, here, we introduce a novel descriptor of HRV, based on a modified Poincaré plot of 24-h RR-recordings. We show that there exists a – regressive biexponential – HRV versus HR “master” curve (“M-curve”) that is highly conserved for a healthy individual on short and medium terms (on the hours to months scale, respectively). At the same time, we reveal how this curve is related to age in the case of healthy people, and establish alterations of the M-curves of heart-attack patients. A stochastic neuron model accounting for the observed phenomena is also elaborated, in order to facilitate physiological interpretation of HRV data. Our novel evaluation procedure applied on the time series of interbeat intervals allows the description of the HRV(HR) function with unprecedented precision. To utilize the full strength of the method, we suggest a 24-hour-long registration period under natural, daily-routine circumstances (i.e., no special measuring conditions are required). By establishing a patient’s M-curve, it is possible to monitor the development of his/her status over an extended period of time. On these grounds, the new method is suggested to be used as a competent tool in future HRV analyses for both clinical and training applications, as well as for everyday health promotion.

## I. INTRODUCTION

As all biological rhythms, heart rate (HR) carries inherent stochastic features [1], usually represented by the beat-to-beat variability of interbeat interval (heart rate variability, HRV). It is generally accepted that HRV is largely influenced by the autonomic nervous system, and, discounting some special cases of arrhythmias which can easily be identified by statistical methods, a positive correlation between HRV and the health state of heart is heuristically established [2]. HRV is a widely used parameter in heart disease characterization, where it is considered to carry an important diagnostic value. An elevated HRV is regarded as the sign of high fitness and adaptability of the heart, while reduced HRV levels are usually associated with various pathological conditions, such as congestive heart failure, diabetic neuropathy, mental disorders, post-traumatic stress syndrome, cancer, etc. [3–6].

Several methods have been introduced for the study of HRV, such as frequency- and time-domain analyses, and nonlinear descriptions. In time-domain analysis, the main descriptors are statistical measures of the variability of beat-to-beat intervals, such as RMSSD, SDNN, SDSD, NN50, etc. [7]. A typical problem considered here is that all these parameters depend on physical activity (and also on HR), so their evaluation over a given time interval will always yield average values [8]. Unless, the patient is examined in a fixed position, the dynamics of his physical activity will inevitably affect the measurement. To solve this problem, various measurement protocols have been elaborated worldwide, amongst others, by the joint European and American task force [9], which often prefer fixed-position registration of the RR time series. This may, in fact, be a solution for the issue, however it is not always feasible, and even if so, there might be significant fluctuations of HR due to other effects (such as excitement due to the examination, etc.), too, that should ideally be ignored when calculating intrinsic HRV.

Other studies, based on a statistical amount of measurements, determine what the optimal range for each HRV parameter is, but usually cannot provide an in-depth analysis for individual cases [7]. As an alternative solution, the well-documented HR-dependency of HRV is taken into account by correcting it for HR, e.g., by normalization with HR [8] or with exp(-HR) [10], which, respectively, assumes an inverse or an exponential relationship between HRV and HR. Fourier components (e.g. LF and HF) derived from fluctuations of the RR(t) curve are used to describe HRV in the frequency range [9]. These are often attributed to sympathetic and parasympathetic nervous system effects, respectively. However, Billman and others pointed out that this assignment is problematic, because both components of the autonomic nervous system actually contribute to both of the LF and HF components [11]. Recently, various non-linear mathematical methods for the description of HRV have become increasingly popular, such as entropy-, detrended fluctuation analysis, Poincaré plots, etc [7]. The latter, for example, has been proven especially useful in detecting certain types of arrhythmias, though, has been less successful in contributing to the general description of the HR-dependence of HRV [12,13].

All in all, without a clear understanding of the HR dependency of HRV, it does not seem possible to find a narrow set of global parameters that would adequately characterize individual persons’ HRV data. The question is whether there could be established a person-specific HRV(HR) function that is clearly defined, and does not explicitly depend on other parameters like time, physical activity and its history, etc., but only on HR. The results of Monfredi et al. imply that, if there exists such a function, it should be of rather exponential than hyperbolic nature [10]. They actually provide a general experiential formula, with a single, decremental exponential, which is apparently characteristic of all mammalian organisms. However, their method of data evaluation, and hence, the standard deviation of their HRV data does not allow its validation for individual cases.

In this paper, we outline an attempt to overcome this obstacle, using a special evaluation method for the HRV time series. Based on a modified Poicaré plot of the data gathered by a wearable heart-rate and activity sensor, we derive a master curve (“M-curve”) for characterizing the HRV(HR) function, that shows remarkable invariance to most other explicit variables (time, physical activity, etc.), and considered to be taken as a specific measure to the individual. If the HR interval is wide enough, the M-curve can normally be fitted with two exponentials, and for more in-depth mathematical description, we introduce a stochastic model on biomimetic grounds. The new analysis is then applied to evaluate a data base containing 24-hour long ECG recordings of healthy volunteers and individuals freshly undergone myocardial infarction. The analysis reveals a statistically significant deviation of model parameters of diseased patients from those in the healthy reference group. Finally, we discuss the potential applications of the new method in various disciplines of clinical science and everyday life.

## II. MATERIALS AND METHODS

The method for the derivation of the “Master curve” describing the HRV(HR) dependence, and its basic features were demonstrated via a case study on a healthy volunteer (39-years-old male), performing daily routine activities. The study was approved by the Ethics Committee of the Medical Research Council (ETT-TUKEB) operating as a board of the Ministry of Human Capacities of Hungary (approval identifier: IV/7109-1/2021/EKU), and conducted according to the WMA declaration of Helsinki. Personal patient information was handled confidentially, and written informed consent was obtained prior to the study. The data of RR intervals were collected by a Polar V800 wearable heart rate monitor, equipped with a physical activity recording feature, validated for scientific use of HRV studies [14]. The evaluation method was then applied to ECG data obtained from the Telemetric- and Holter-ECG Warehouse (THEW) at the University of Rochester Medical Center, New York, United States [15]. 24-h Holter recording data of 202 healthy volunteers (Database Normal, EHOL-03-0202-003, age ranging from 9 to 82 years) and 93 patients with acute myocardial infarction (Database AMI, E-HOL-03-0160-001, age ranging from 27 to 90 years) were analyzed. The time series were filtered for outliers by a sequential cluster analysis using the “dbscan” routine of MATLAB (MathWorks, 2020).

## III. RESULTS AND DISCUSSION

### A. DERIVATION OF THE M-CURVE

For the characterization of the point-by-point HRV, the Poincaré plot is the most popular tool [12,13]. Fig.1a shows the traditional Poincaré-representation. Since these plots are quasi-symmetric to a line making 45 degrees with the X and Y-axes, a Poincaré analysis of the RR or HR time series often involves fitting of a tilted ellipsis to the plot, in order to characterize the extent (the “width” and “length”) of the set of points depicted, with respect to the symmetry axis. Accordingly, such an evaluation describes the time series with two numbers, corresponding to the maximal RMS of HRV, and the span of HR values [13]. In order to reveal a more detailed dependence of HRV from HR, we started from a modified Poincaré-plot, defined by the following equation:

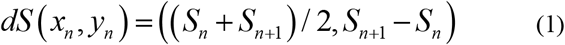

where S may stand for RR or HR, and d for difference. A notable feature of this representation is that now the set of points is quasi-symmetric to the X-axis, which may be utilized in the analysis of other time series, as well. (Note that (1) is formally similar to the renowned Bland-Altman plot, often used for comparison of two time series, however, here we use subsequent points of the same time series, instead [16].) For the sake of convenience, we chose the following version of (1):

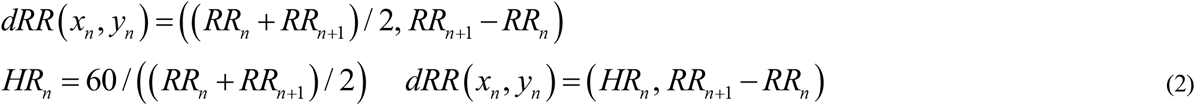

i.e., we practically described the point-by-point difference of subsequent RR values as a function of mean RR (Fig. 1b) and HR (Fig.1c). If there are too many points to distinguish on the plot, one may color-code for point density, to better describe their distribution (Fig.1c). Instead of fitting an ellipsis to the point set, we determined the RMS of the dRR values for each HR (the latter defined by 2/(RR_n_+RR_n+1_)), using the symmetric feature of the point distribution to the X axis. This treatment allowed the determination of a characteristic HRV parameter as a function of HR, with a higher precision than earlier attempts (Fig. 1d). (For a more detailed discussion of differences and similarities with previous treatments, see Supporting Information.)

**FIGURE 1.**
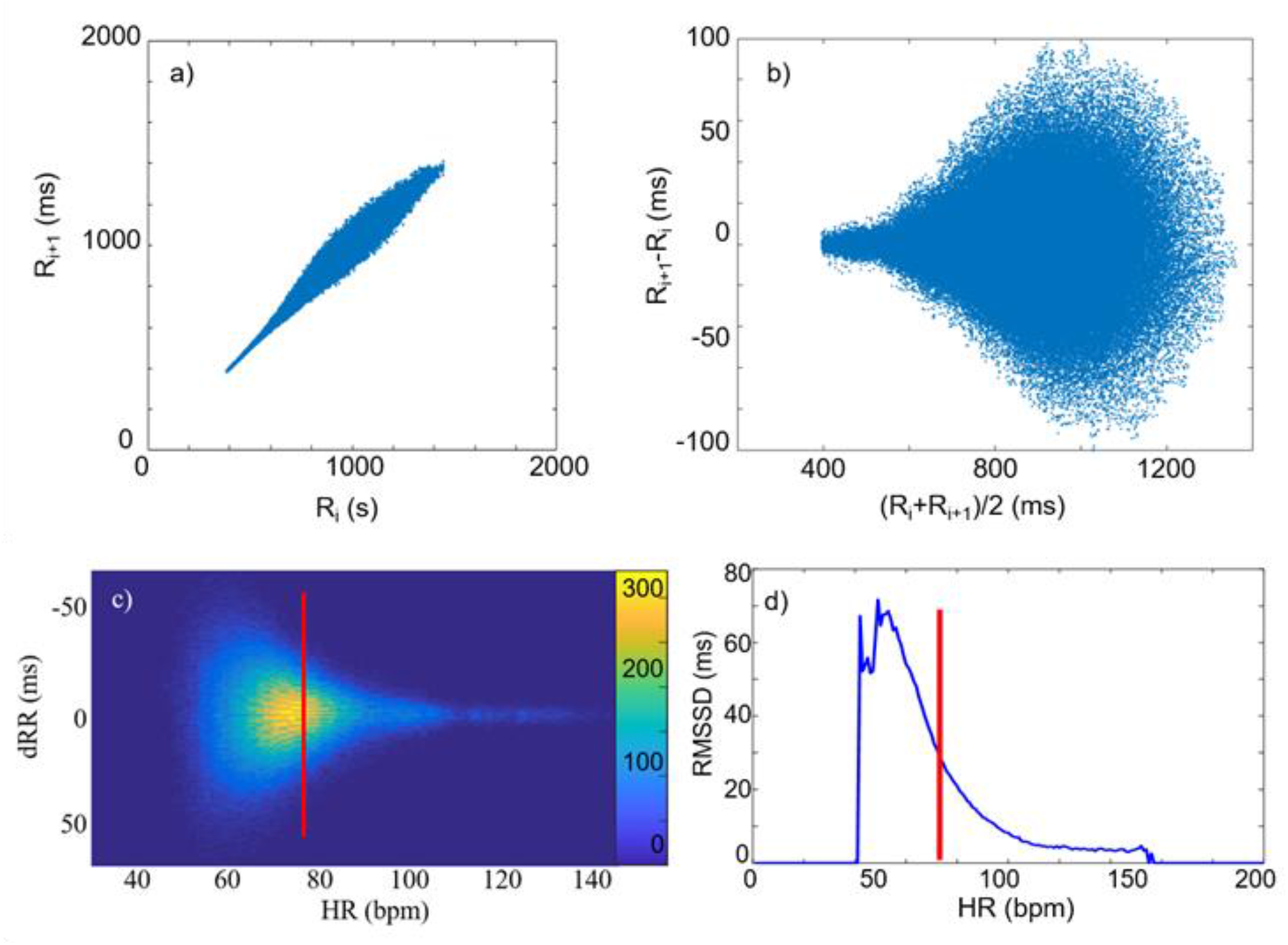
(a) Poincaré plot of a typical RR time series. (b) The same in Bland-Altman-like representation. (c) dRR as a function of HR, as calculated from the data in (b). The color code is to show the frequency of the data. (d) The RMSSD versus HR curve, as calculated from data in (c).

In the following, first we demonstrate through a case study that this function shows remarkable conservation features, and it is specific to the subject (Master curve or “M-curve”). Next, we apply the evaluation method to analyze RR-data of healthy and diseased individuals. Finally, we establish a stochastic model to formally describe the M-curve, and hypothetically associate the parameters of the model to some physiological descriptors of the autonomic nervous system.

### B. INDEPENDENCE OF THE M-CURVE FROM ACTIVITY AND DATE

Fig.2a shows the M-curves determined before, during and after a 1-hour long sub-maximal training of a volunteer. It can be seen that in the common HR range, the two curves are overlapping each other, i.e., the M-curve follows the same trend before and after training. It is often established in the literature that the RR variability decreases during, and shortly after, a physical exertion [17]. Our analysis reveals that, on the hours scale, this effect does not accompany with a change of HRV at a certain HR, but rather with a “shift” on the M-curve towards the higher-HR region reached during, and shortly after, the exercise. (Note, however, that these results do not exclude deviations from the M-curve on a shorter time scale, e.g., that of minutes.) Apparently, different activities related to the daily routines on different days do not influence the M-curve of an individual on a daily basis, either. According to our measurements, though, these factors may well change the HR range, but the actual shape of the 24-hour M-curve will not be effected.

**FIGURE 2.**
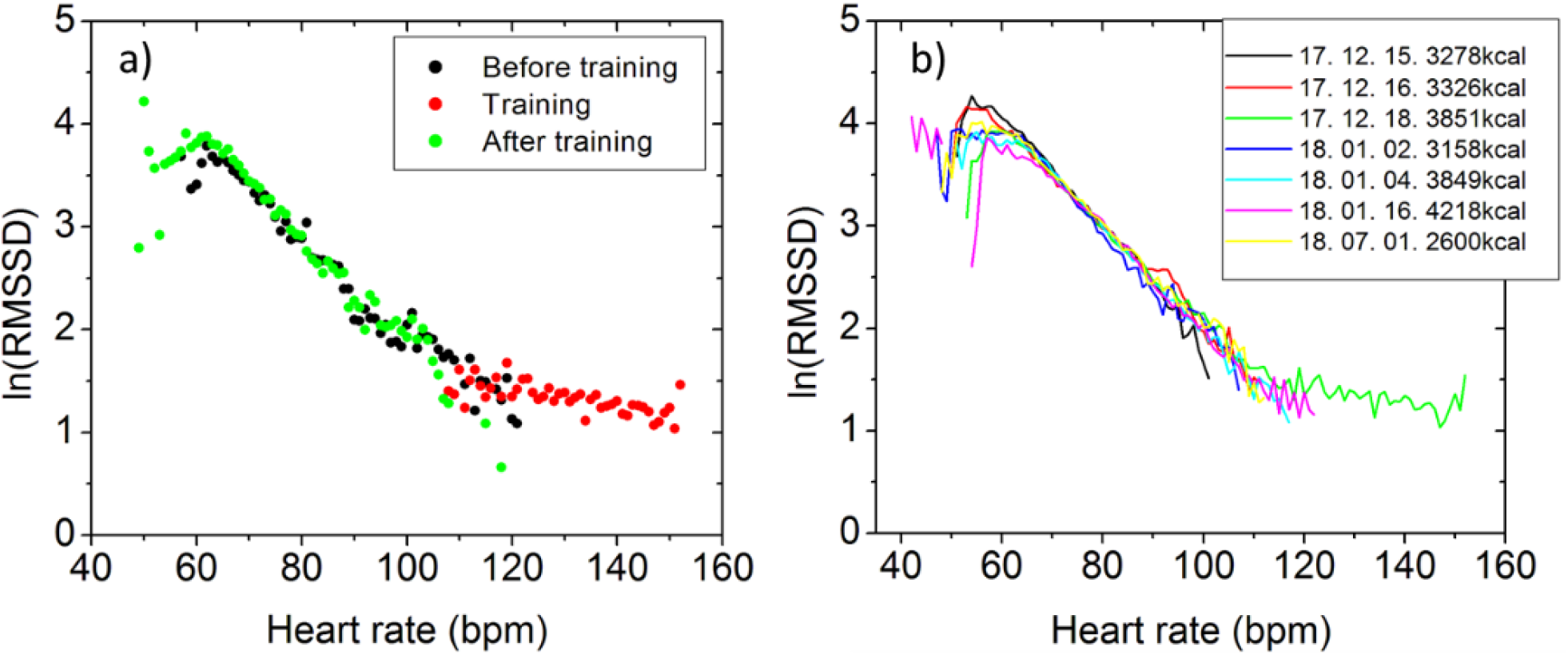
Reproducibility of M-curve. (a) M-curves calculated from data recorded before, during and after a submaximal training of a volunteer (<85% of max HR). (b) M-curves recorded on individual days. The insert shows the dates of recording and the corresponding cumulative physical activities in kcal.

The question arises, whether or not the reproducibility of M-curve persists on longer time scales, too. For this reason, we registered RR data of the same volunteer on subsequent days or weeks, and after a half-year-long intermission period. Fig.2b shows a series of daytime M-curves calculated from the data collected. Although, the daily activities (shown in the insert) are very different (due, e.g., to occasional trainings), the central part of the M-curves remained practically the same. Significant differences occur only at the extrema of HR, i.e., the maximum and minimum HR values of the curves show variation, due, e.g., to the presence or absence of a strong training on the particular day that seem to shift the HR set towards higher values. (The steep cutoffs at small HR values are an artifact of the method due to sparse data.)

### C. GENERALIZATION AND AGE DEPENDENCE

Since all the above findings were demonstrated via a case study, the question arises whether similar statements apply to other individuals, too. Processing data from 202 volunteers and additional 93 patients in the THEW database, we found that the M-curves of the vast majority (> 95%) of healthy volunteers and a considerable part (> 50%) of diseased patients showed the same, biexpoential-type pattern. It should be noted, however, that the “second” (slower-changing) exponential-like component was present only in those cases where the recorded HR range was sufficiently wide (that is, when the HR extended ca. 110 bpm). For different people, the M-curves showed smaller or greater differences, contrary to the location of the transition range, which appeared to be rather conservative, between 110 and 130 bpm.

In order to reveal any systematic dependence of the HRV on the age of the healthy individuals, we performed an age-class cohort study to determine the “averaged M-curves” for several age groups (Fig.3a). In other words, we calculated the mean HRV values of individuals belonging to each age group, at each HR value. Notably, these group-averaged M-curves show the characteristic feature of two exponential-like phases.

**FIGURE 3.**
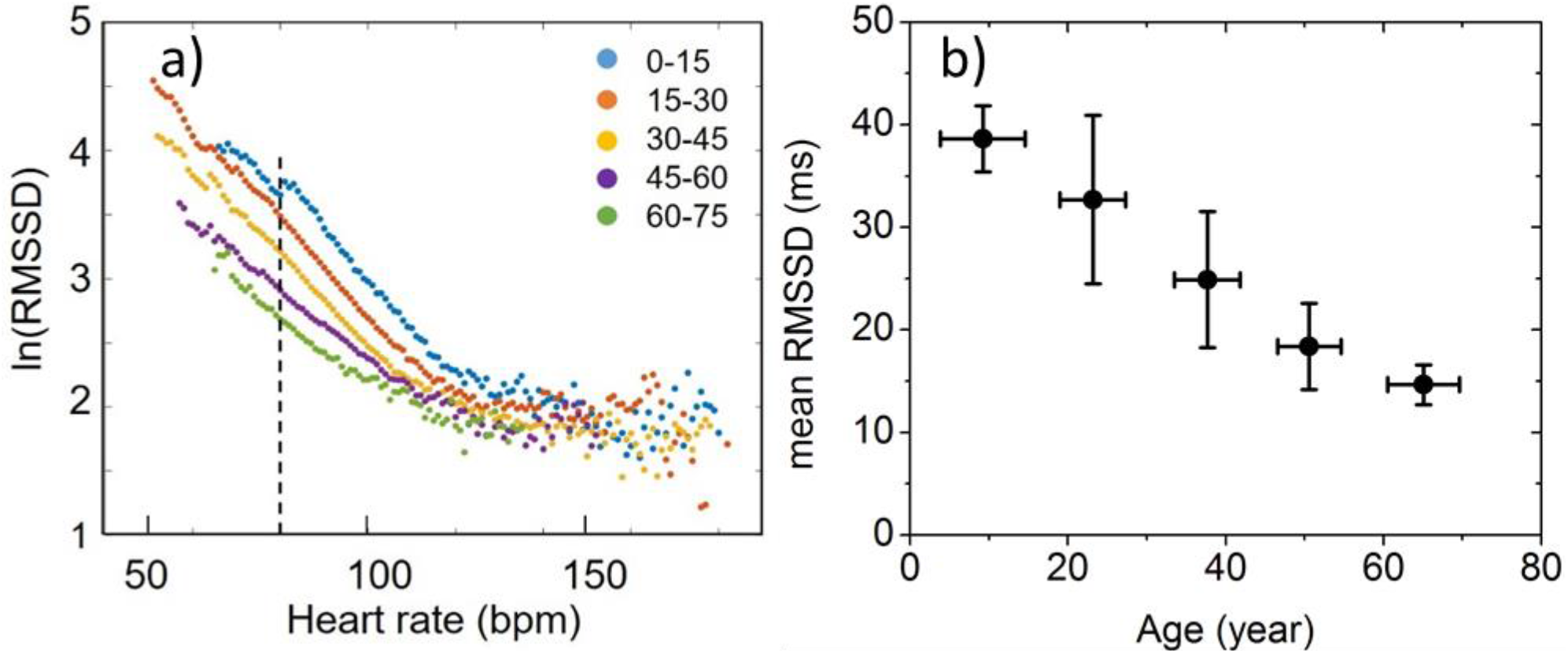
(a) Group-averaged M-curves of healthy volunteers belonging to different age classes. Sections with the dashed line identify the mean RMSSD values at HR=80. (b) Cohort mean HRV(80) values as a function of age.

According to our analysis, the, e.g., HRV(80) data give a decent measure to distinguish HRV data upon age change (Table 1, Fig.3b). Note that similar statements concerning the age-dependence of HRV have been established earlier on different grounds [18]. In the most comprehensive study, Tsuji et al. determined the SDNN of interbeat intervals and the average heart rate from 2-hours-long ECG recordings of each patient. Investigating age-selected cohorts of 1192 healthy subjects, they found that HRV was determined by age and HR to a different extent, but both in an “inversely associated” manner.

**TABLE I.**
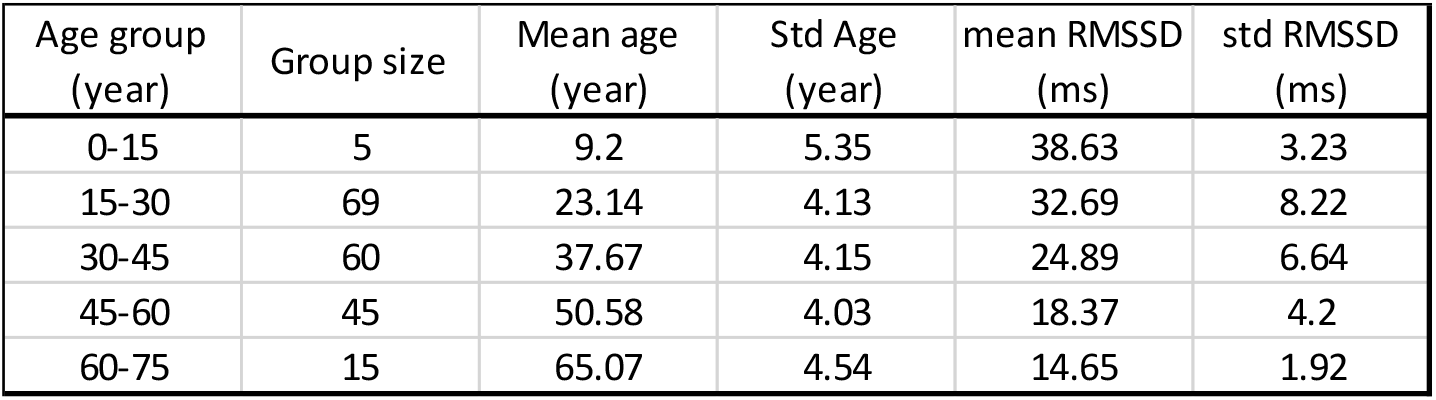
Age-group distribution and the corresponding HRV parameters of healthy individuals of the database analysis shown in Fig.3.

| Age group (year) | Group size | Mean age (year) | Std Age (year) | mean RMSSD (ms) | std RMSSD (ms) |
| --- | --- | --- | --- | --- | --- |
| 0-15 | 5 | 9.2 | 5.35 | 38.63 | 3.23 |
| 15-30 | 69 | 23.14 | 4.13 | 32.69 | 8.22 |
| 30-45 | 60 | 37.67 | 4.15 | 24.89 | 6.64 |
| 45-60 | 45 | 50.58 | 4.03 | 18.37 | 4.2 |
| 60-75 | 15 | 65.07 | 4.54 | 14.65 | 1.92 |

### D. EFFECT OF DISEASE

To see whether heart disease may have an effect on the M-curve, we applied the method to data of patients with acute myocardial infarction (AMI). In this case, we identified a number of unusual patterns on the modified Poicaré-plots (e.g, Fig.4a) indicative of different types of arrhythmias [13], hence, the corresponding M-curves appeared in various anomalous shapes (data not shown). Nevertheless, for some 50% of the AMI patients, the M curves showed the “regular” biphasic decay. Considering only these data, we calculated the averaged M-curves of healthy and hospitalized AMI patients belonging to the same age group (between 45 and 60 years, 45 and 55 patients, respectively), shown in Fig 4b. It can be seen that the latter curve runs below the one of the control group in the 60 < bpm < 100 HR range.

**FIGURE 4.**
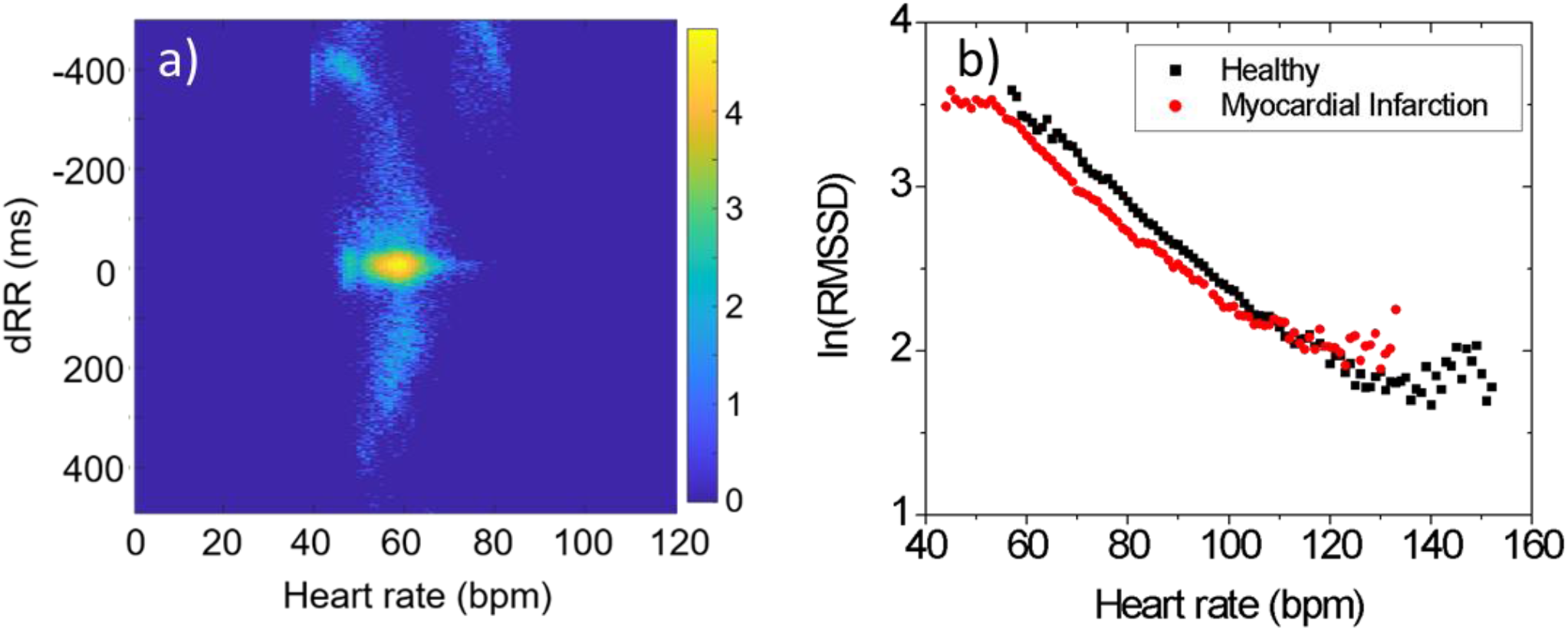
HRV(HR) characteristics of patients with myocardial infarction. (a) Bland-Altman-type representation of HRV data of an individual typical of arrhythmia. (b) Cohort-averaged M-curves of healthy and heart-attack patients (black and red symbols, respectively), belonging to the same age class (45-60 years).

### E. COMPARISON TO EARLIER RESULTS ON THE HRV(HR) DEPENDENCE

Most of the traditional time-domain analyses of HRV usually yield one (or a few) global parameter(s) that characterize the variability of the whole HR (or RR) time series. This is normally a sort of averaged HRV value, corresponding to the HR time series registered under pre-defined particular measuring conditions, this way limiting the HR range, in order to avoid the HR-dependence of HRV.

Since the high-profile publication of Monfredi et al. [10], there is an ongoing debate about the “proper normalization” of HRV by HR, as well as about the possible “disentanglement” of the two state-describing parameters [17]. According to the, perhaps, most accepted approach, the actual HRV measure should be normalized by the average HR [8,9,19,20] correcting for the well-known fact of HR-dependence of HRV, assuming a reciprocal relationship between the two. This looks a logical method if one assumes that the distribution of HRV, measured normally as RMSSD or SDNN, is constant in the time domain, where the registered RR time series are naturally represented. This would also imply that HRV could be described by a single parameter, which is an appealing perspective. On the contrary, Boyett et al. claim that the HRV(HR) function is a single, unique exponential for (healthy) humans and mammals, in general. On the one hand, this would mean a steeper dependence than the simple reciprocal relation, implying that data derived using the latter assumption are flawed, and on the other, would establish a strict coupling between HRV and HR, implying that the HRV-effects are, mostly and simply, due to the change of HR, impairing the diagnostic value of the former. They constructed a simpler and a more complex biophysical model based on the stochastic nature of ionic currents charging the membrane of the pacemaker cells, to interpret the exponential-like relationship between HRV and HR they inferred. Although, both models were able to account for describing the descending tendency of the HRV data as a function of HR, the fit of the simulated values to the experimental ones showing fairly high standard deviation, was rather poor in both cases [10].

VanRoon et al. argue that the quality of the gathered data does not allow to assess their exponential or reciprocal dependencies on the HR scale spanned, but votes for the simpler (reciprocal) case [21]. Gasior et al. use an improved correction factor proportional to the inverse 3^rd^ power of HR (see also Supporting Information) [22]. In a recent paper, vandenBerg et al. investigated 10-s ECG records of a population of 13,943 individuals, both males and females, with a wide age distribution, and tested 4 different methods for correcting HRV by HR, assuming linear, exponential, hyperbolic or parabolic relationship between the two quantities. They also concluded that the data scattered too much to allow clear distinction among the 4 cases, however they found the exponential correction slightly superior to the others, though, not perfect, especially at high HR values [23]. In view of this debate, our low-noise M-curves can be decisive. The linearity of the M-curve in semi-logarithmic plot clearly shows an exponential type of dependence at HR values below ca. 100 bpm. On the other hand, we also show that the general HRV(HR) function can be described by two components, whose slopes are usually different for different subjects (see below), so the M-curve may be taken as characteristic to the individual, and it is rather conservative on the time scale of days or even months. On the scale of decades, however, the M-curve does depend on age, indicating an overall decrease of HRV for elderly people, and a similar effect can be observed for patients of heart disease. Our results also imply that it is generally not enough to characterize HRV by a single parameter (e.g., after proper normalization of the raw data), but the whole HRV(HR) function must carry important physiological information. In order to facilitate its deeper interpretation, we elaborated a stochastic model that can account for the characteristic features of the M-curve.

### F. STOCHASTIC MODEL

Following earlier approaches [10, 24], we considered a simple integrate-and-fire neuron model to mimic HRV, where a white noise (δ) of Gaussian amplitude distribution of “d” RMS is added to the charging ion current (I_t_) (Fig.5).

**FIGURE 5.**
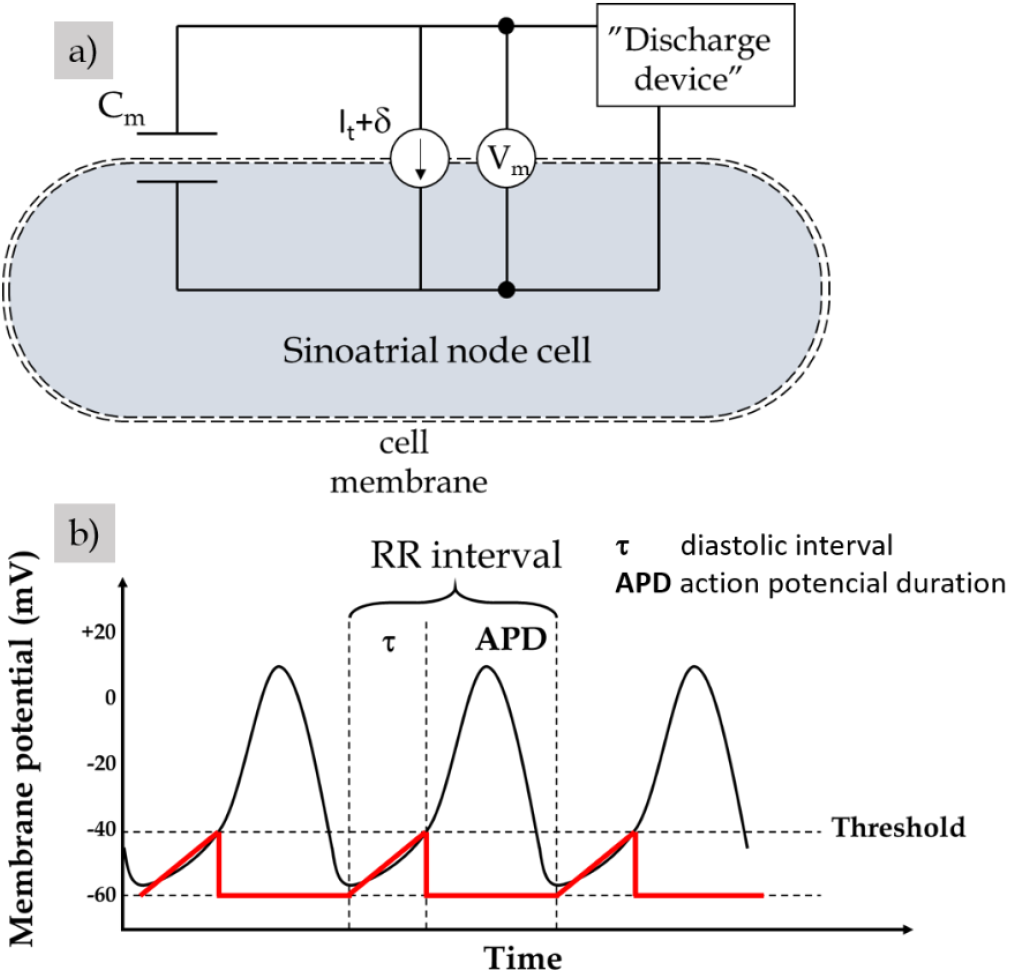
A simple, stochastic integrate-and-fire model of HR and HRV. (a) An electrical substitution circuit of the model. C_m_ is the membrane capacitance, and V_m_ is the membrane potential which is short-cut by a discharge device, if it exceeds a threshold level. (b) Schematic representation of the time course of V_m_. Black line symbolizes the physiological signal, while the red line is the outcome of our model.

Due to the presence of the noise term, the time intervals between adjacent firing spikes (representing the RR intervals in this model) show a stochastic distribution even at constant I_t_. It is, however, assumed that I_t_ >> d, so HR is considered to scale with I_t_.

In mathematical terms,

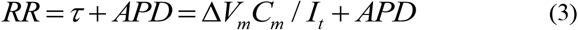

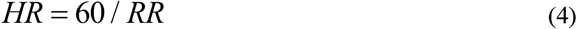

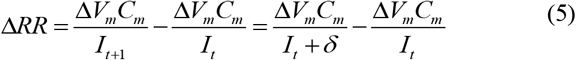

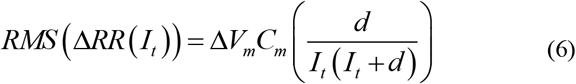

where δ is the instantaneous noise term, “d” denotes RMS(δ), ΔV_m_ is the difference between the negative peak and the threshold, C_m_ is the membrane capacitance, APD is the action-potential duration (considered to be 160 ms), and τ is the diastolic interval [10]. If one assumes a constant “d”, as in [10], it is easy to see from the equations, that one gets back an approximate hyperbolic relationship between HRV and HR, which, as we discussed above, cannot fit the measured data with decent precision. Hence, we allowed a variation in “d” as a function of I_t_, and determined the d(HR) function by fitting the model to the experimental data (i.e., to the M-curve).

As it can be seen in Fig.6a, the d(HR) function can be decently approximated by two linear functions of positive (α) and negative slopes (-β), respectively, below and above a transition range around 110 bpm.

**FIGURE 6.**
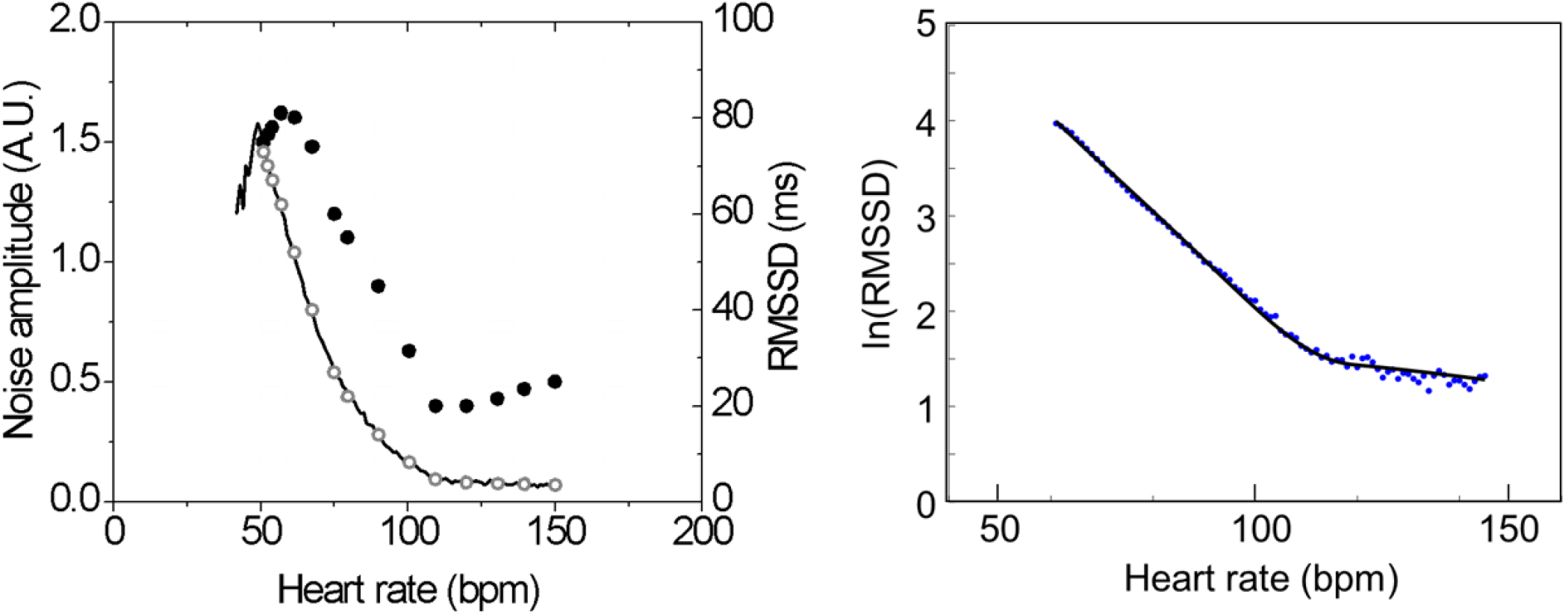
(a) Experimentally determined RMSSD values (open circles) and the result of model-fitting (solid line), as a function of HR. The corresponding noise amplitude (“d” parameter) of the model, as determined from (6). (b) The result of model-fitting of the M-curve in semi-logarithmic representation.

In mathematical terms, the instantaneous δ(I_t_) terms can be represented as:

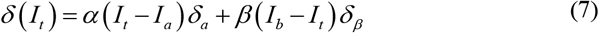

where δ_α_ and δ_β_ are stochastic multiplicators sampled from independent normal distributions of 0 mean and 1 standard deviation. From this, “d(I_t_)” can be expressed as follows:

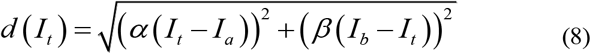

Fitting the experimental M-curve by the set of equations (3), (4), (6) and (8), we could establish that the model is able to describe the experimental data with high accuracy (Fig.6b), and the α, β, I_a_ and I_b_ parameters can be determined (see also SI). Based on this result, it is straightforward to assume that the noise term of our model (“d”) can be considered as a result of two stochastic processes dominating below and above the transition zone. While the contribution of the former one shows a descending tendency with increasing HR, the weight of the latter one is slightly increasing with it.

Without aiming to give a strict physiological interpretation of the parameters of the stochastic model able to describe the experimental M-curve, we cannot resist calling the attention to the striking similarity between their behavior and that of the components of the autonomic nervous system. According to the widely accepted view, HR is determined by the sympatho-vagal balance, in which framework increasing parasympathetic activity decreases HR and increases HRV, while increasing sympathetic activity acts oppositely (Fig.7). In the absence of vegetative control (e.g., during autonomic blockade), an intrinsic HR is set around 100-110 bpm, below which value parasympathetic, while above it sympathetic effects dominate. Given the same tendency established for the noise components of our model, it is reasonable to assume a close connection between these and the components of the ANS, however, a more established support for this hypothesis should be the subject of follow-up studies.

### G. POSSIBLE APPLICATIONS OF THE M-CURVE

Even in the absence of a solid physiological interpretation of the data, knowing the tendency of the characteristic statistical parameters (e.g., HR(80), α and β in Fig.S3) as a function of age, one can establish reference values for the age groups. Hence, we suggest a protocol for HRV registration and evaluation based on the new method: A preferably 24-hour (or longer) RR-recording should be registered once a year, from which the M-curve characteristic of the patient can be determined by high accuracy. In between, shorter measurements - still appropriate to determine the HR(80) value by high precision - are sufficient to follow changes in the status.

**FIGURE 6.**
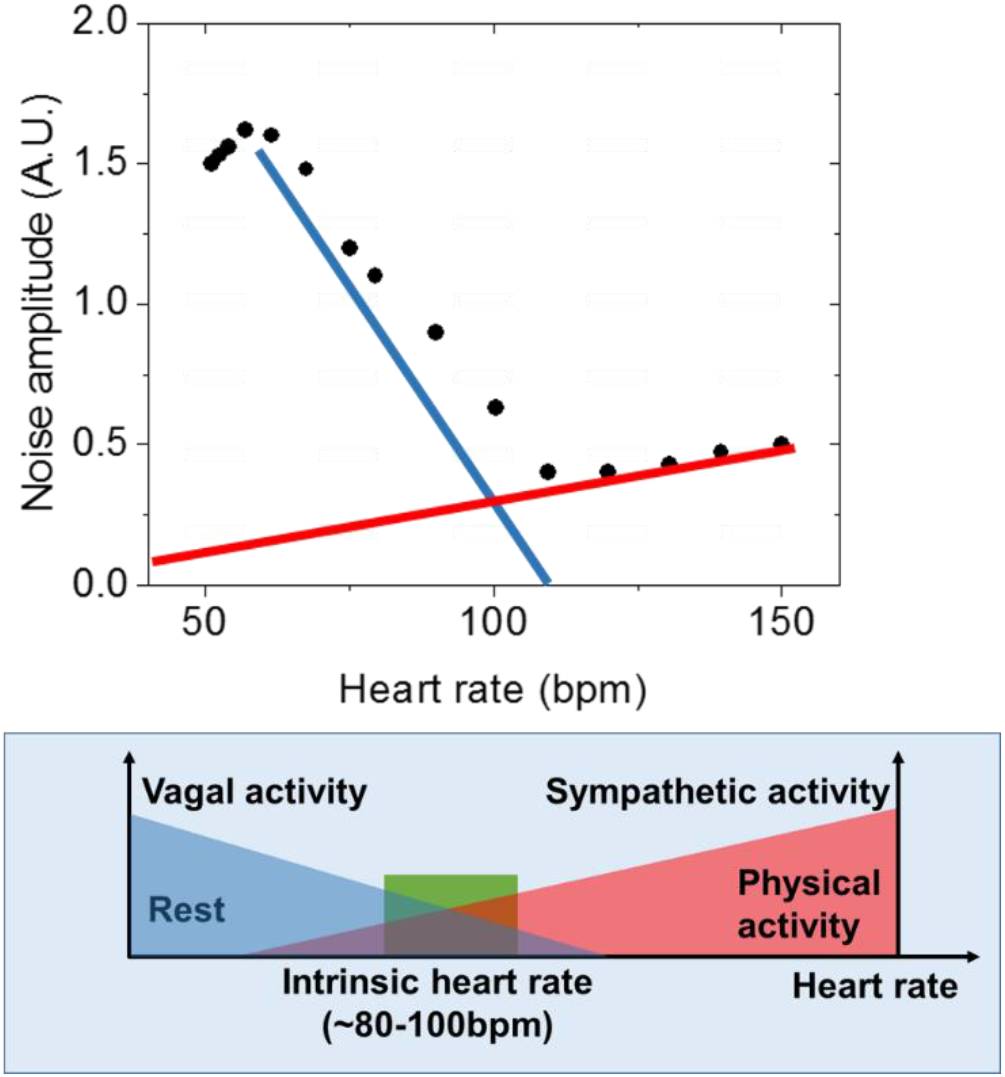
Similarity between the HR-dependences of the two components of our noise parameter (d) and the two components of the autonomous nervous system (ANS), as generally assumed [25].

In addition to the opportunity of medium and long-term monitoring, the precise description of the M-curve allows the determination of an instantaneous HRV value on the minutes scale, that is supposed to be characteristic to the momentary state of the patient, independent from the actual HR value. For this purpose, the exact knowledge of a person’sHRV (HR) function would be essential, but the presently applied methods either use a raw measure of HRV, or correct it by an ill-defined normalization function (see Figure S2). Since the M-curve describes the medium-term HRV(HR) function with high precision, normalization of a short-term (e.g., 5-minute) HRV(HR) recording according to the M-curve should be informative for the actual state.

## IV. CONCLUSIONS

We introduced a new representation of heart rate variability data, based on a Bland-Altman-like mathematical transformation of the Poincaré-plot, that allows a natural visualization of the beat-to-beat variability of interbeat intervals as a function of heart rate. The graphs of an individual show striking reproducibility on the daily and monthly scales, and physical activity also does not seem to affect their shape, only causes shifts along the same curve (that we call Master- or M-curve). Recordings of beat-to beat intervals on the hours scale allow the construction of a high-quality M-curve, determining the HRV(HR) function with unprecedented precision, as compared to the conventional representations.

As a function of HR, in a semi-logarithmic plot M-curve shows a linear dependence with negative slope between ca. 60 and 100 bpm, while above this interval the rate of regression is weaker, implying a biexponential-like decay. Analyzing data from a public ECG-database, we found that the M-curves of the vast majority (> 95%) of healthy volunteers and a considerable part (> 50%) of diseased patients showed a similar pattern. For different people, the M-curves showed smaller or greater differences, but arranging the data in age groups revealed a clear shift of the averaged M-curves to lower ranges, by progressing age. The HRV(80) data are a good representation of this tendency, and the averaged values can be taken as a normal reference. The plots of heart patients with myocardial infarction differ significantly in shape and/or range.

Nevertheless, our results also suggest that a single parameter is not sufficient to fully describe the complex features of HRV data. We show that an integrate-and-fire stochastic neuron model is able to fit the experimental HRV(HR) graphs of healthy people with high precision, at the same time offering plausible clues for a physiological interpretation of the HRV(HR) relationship.

The new method is suggested to be used as a competent tool in future HRV analyses, for both clinical and training applications, as well as for everyday health promotion (in cardiology, polysomnography, sports, training, etc.).

## Supporting information

### S1. Differences and similarities to the conventional evaluation methods

In a conventional time-domain analysis, one could determine the HRV(HR) dependence by a sectional evaluation of the HR time series, for a rolling time window (similarly to Monfredi et al.) [10]. However, when selecting the optimal width of the time window, one has to consider the problem stemming from the time-dependence of HR. If the analysis is restricted to short time intervals (where HR does not change significantly), it results in a higher uncertainty in HRV, while longer averages accompany with information loss on the HR scale (Fig.S1). Our evaluation method seems to solve this problem.

**Figure S1.**
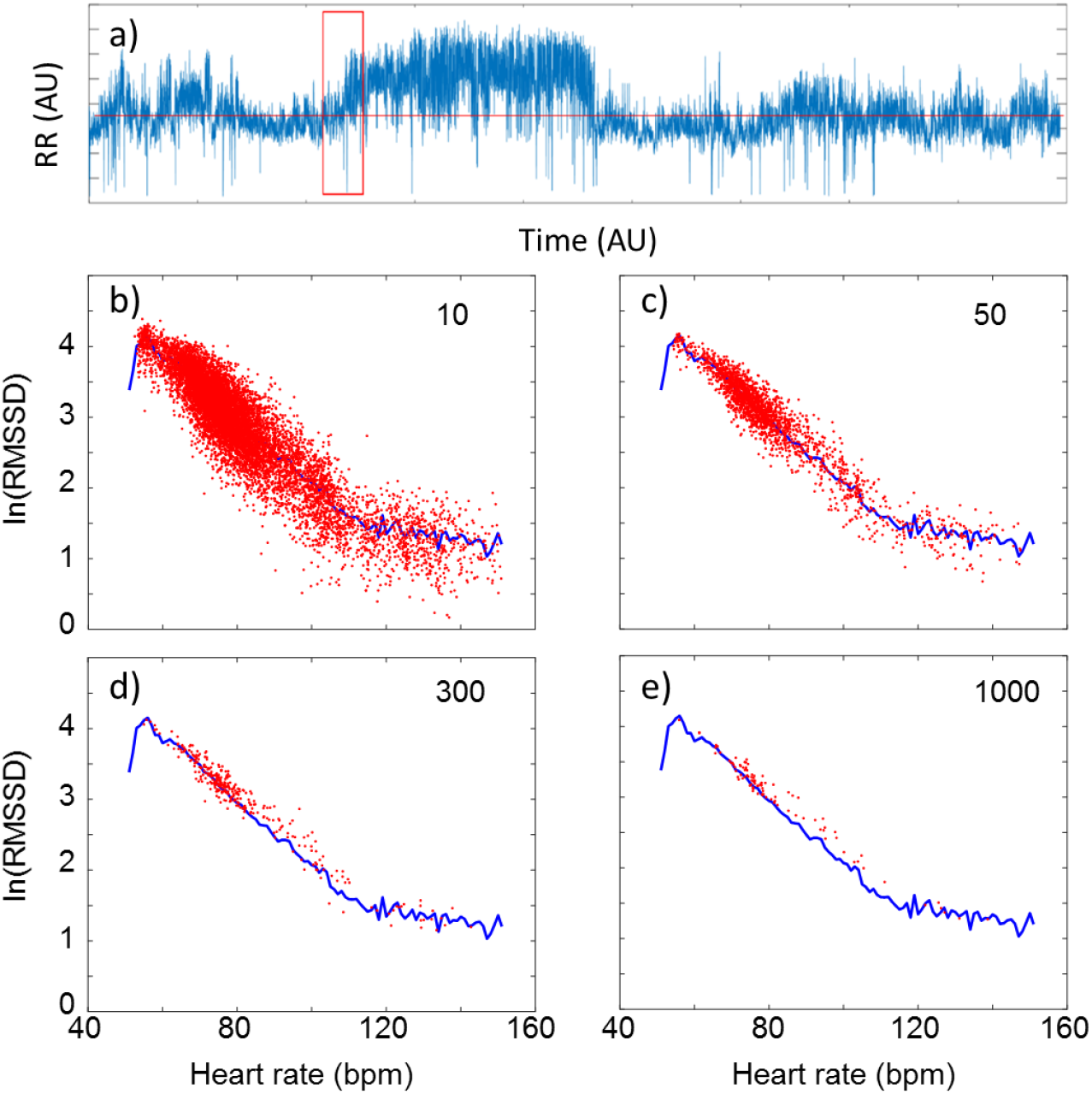
Comparison of the M-curve (blue line) with data of the conventional moving average evaluation (red points) in semi-logarithmic representation. a) An example RR time series with the moving window. (b), (c), (d), (e): Data obtained with window size of 10, 50, 300 and 1000 data points, respectively.

In a semi-logarithmic representation, the core part of the M-curve (between ca. 60 and 100 bpm) shows a linear dependence with negative slope, while at around 100 bpm, there is a transition range, above which the rate of regression is weaker in the function of HR. (Note that the latter component can only be seen if HR extends well beyond this transition range or “break point”.) This means that in the full HR range, the HRV(HR) function follows a regressive exponential-like dependence of two decaying components.

### S2. Normalization attempt of HRV by HR

Establishing a single HRV parameter characteristic to the cardiac health state of the patient has always been an appealing goal. Hence, a proper normalization function that could account for the observed HR-dependence of time-domain HRV data has been intensively searched for. In the main body of the paper, we discuss that, apparently, a universal analytical function cannot be given to describe the HR-dependence of HRV of each individual in the “full” HR-range. So far, however, the quality of HRV data obtained by earlier methods, has not allowed to make such a definite statement, leaving room for a lot of debate [8,10,17,21,22]. Some clever correction functions, such as using the inverse cubic normalization by HR [22], e.g., seem to give a decent result for moderate HR values, if HRV data are established by the moving-average evaluation. A more accurate representation of the HRV(HR) function (i.e., the M-curve), however, reveals the limitations of this method (Figure S2).

**Figure S2.**
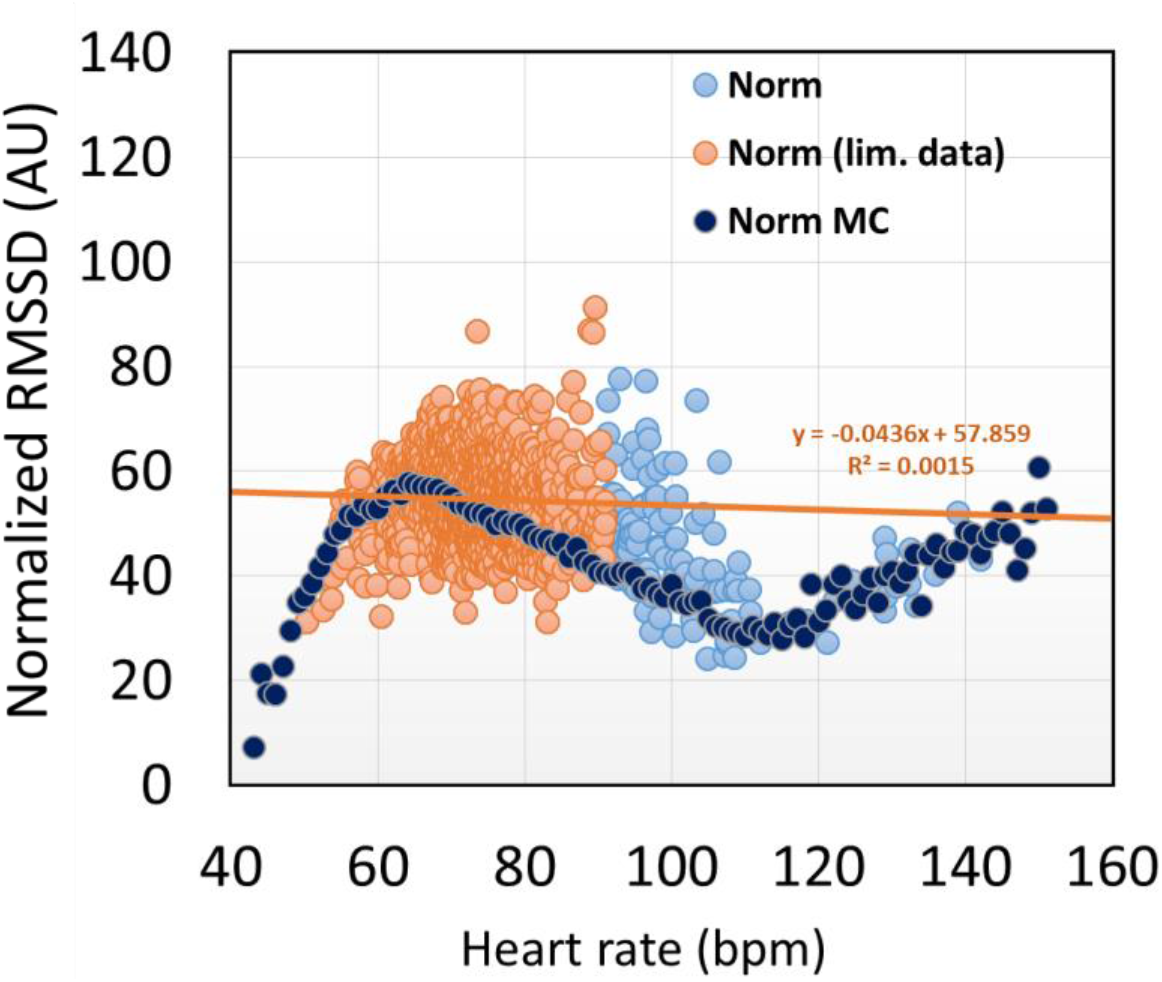
RMSSD data of a typical HR time series normalized by a HR-3 function. Red and light blue symbols: data from a moving-average evaluation, dark blue: data from the “M-curve” evaluation. A decent fit is got to the red-symbol region, unlike to the more accurate data.

### S3. Model fitting to measured data

M-curves calculated from data of healthy individuals could be fitted by our integrate-and-fire model with decent accuracy, however, the standard deviation of the fitting parameters (α, β, Ia and Ib) was rather high (see Fig. S4 a-d, where HRs and HRp were determined from Ia and Ib, respectively, according to Eqs. (3) and (4)). The primary reason for this was the limited HR range of recordings in a number of cases, not allowing the accurate determination of all these parameters. Since only β showed a significant tendency as a function of age, and the other fitting parameters were fluctuating around virtually age-independent average values, we repeated the fits with HRs and HRp values fixed at the mean of their distribution. This restriction did not considerably affect the goodness of individual fits, and worked properly for the group-averaged M-curves, too (Fig. S3). Fig. 8b shows the age dependence of α and β parameters for the latter case. It is apparent that β shows a significant monotonic decrease by age, while the change in α is minor.

**Figure S3.**
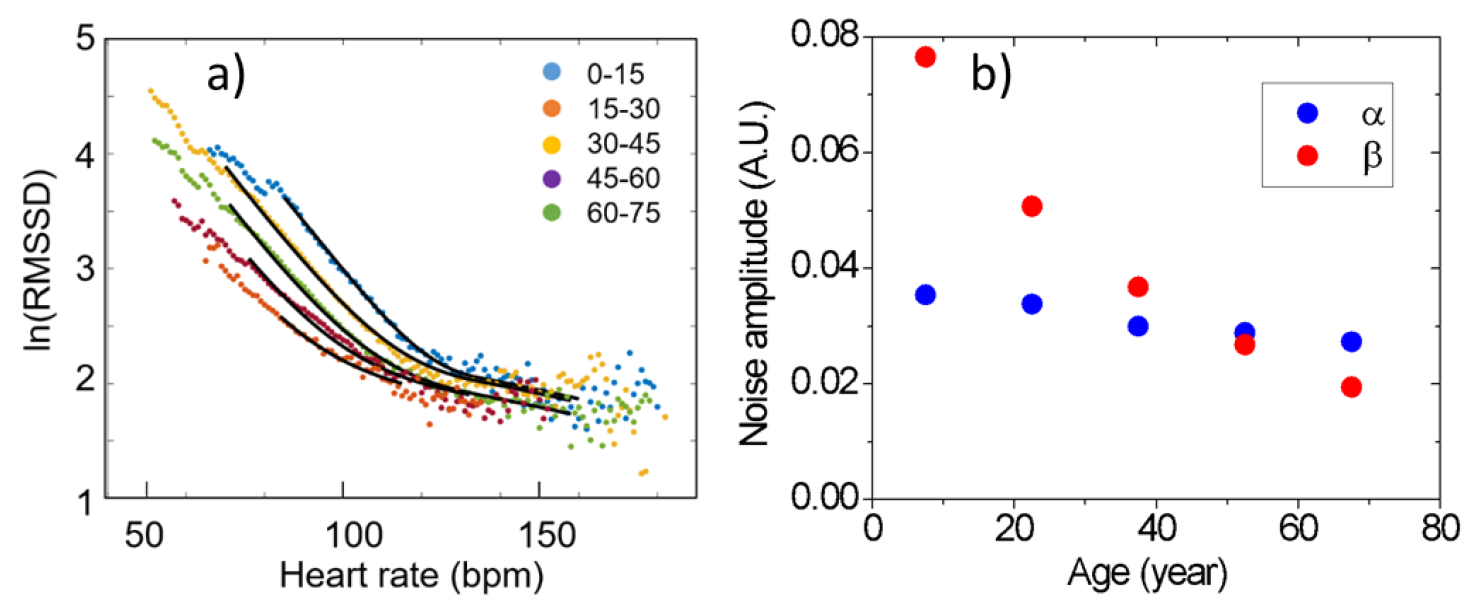
Model fitting of cohort HRV data of healthy individuals from the THEW database [15]. (a) Averaged M-curves of age groups distinguished by color code, and the fitted curves (black lines). (b) Age-dependence of the α and β parameters, while HRp and HRs values kept fixed.

**Figure S4.**
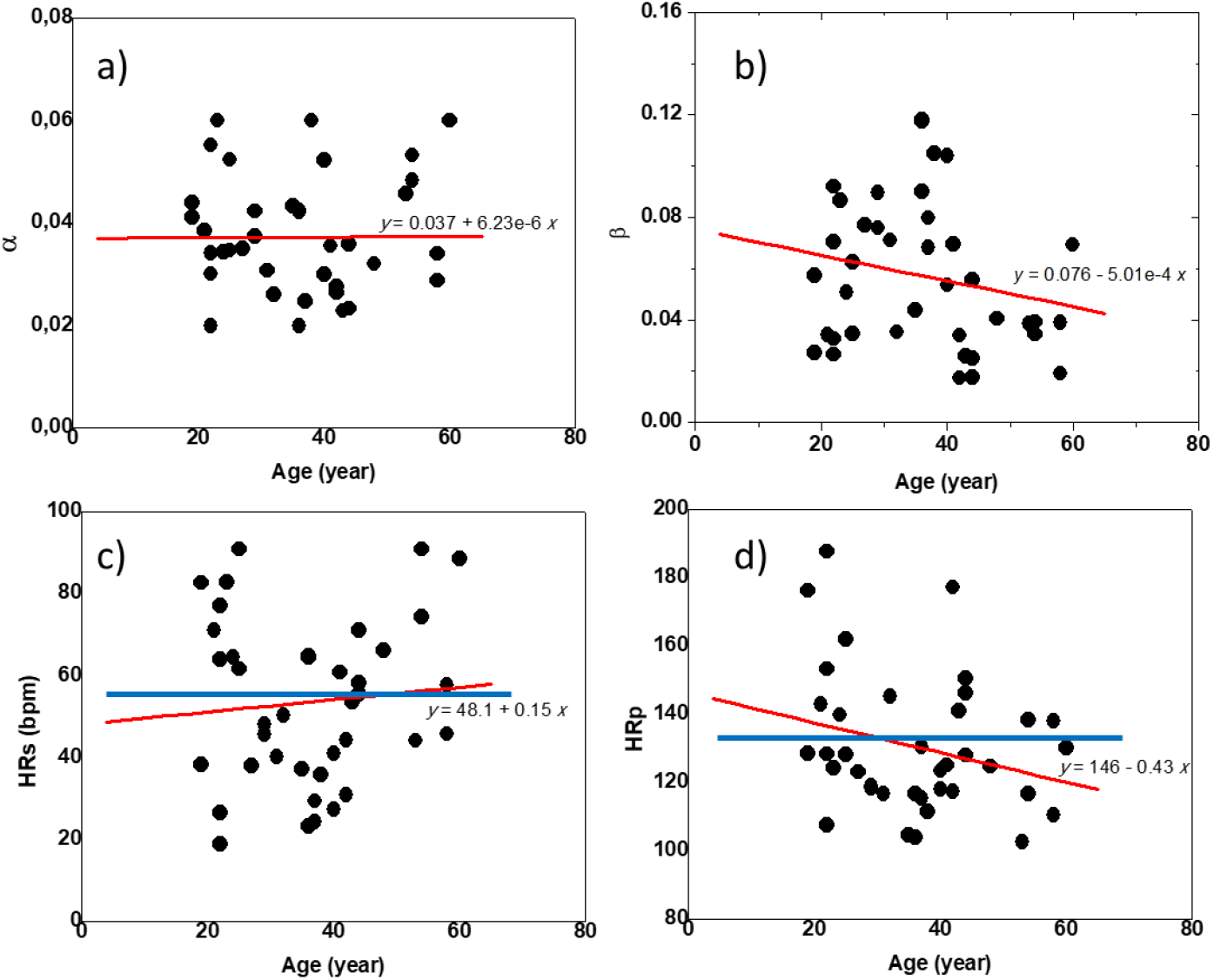
Variable parameters of model fitting of M-curves of healthy individuals from the THEW database [15]. (a) α, (b) β, (c) HRs, (d) HRp.

